# Impact of Influent Carbon to Phosphorus Ratio on Performance and Phenotypic Dynamics in Enhanced Biological Phosphorus Removal (EBPR) System - Insights into Carbon Distribution, Intracellular Polymer Stoichiometry and Pathways Shifts

**DOI:** 10.1101/671081

**Authors:** Nehreen Majed, April Z. Gu

## Abstract

This study investigated the impact of influent carbon to phosphorus (P) ratio on the variation in P-removal performance and associated intracellular polymers dynamics in key functionally relevant microbial populations, namely, PAOs and GAOs, at both individual and populations levels, in laboratory scale sequencing batch reactor-EBPR systems. Significant variations and dynamics were evidenced for the formation, utilization and stoichiometry of intracellular polymers, namely polyphosphate, glycogen and Polyhydroxyalkanoates in PAOs and GAOs in the EBPR systems that were operated with influent C/P ranged from 20 to 50, presumably as results of phylogenetic diversity changes and, or metabolic functions shifts in these two populations at different influent C/P ratios. Single cell Raman micro-spectroscopy enabled quantification of differentiated polymer inclusion levels in PAOs and GAOs and, showed that as the influent rbCOD/P ratio increases, the excessive carbon beyond stoichiometric requirement for PAOs would be diverted into GAOs. Our results also evidenced that when condition becomes more P limiting at higher rbCOD/P ratios, both energy and reducing power generation required for acetate uptake and PHB formation might shift from relying on both polyP hydrolysis and glycolysis pathway, to more enhancement and dependence on glycolysis in addition to partial/reverse TCA cycle. These findings provided new insights into the metabolic elasticity of PAOs and GAOs and their population-level parameters for mechanistic EBPR modeling. This study also demonstrated the potential of application of single cell Raman micro-spectroscopy method as a powerful tool for studying phenotypic dynamics in ecological systems such as EBPR.

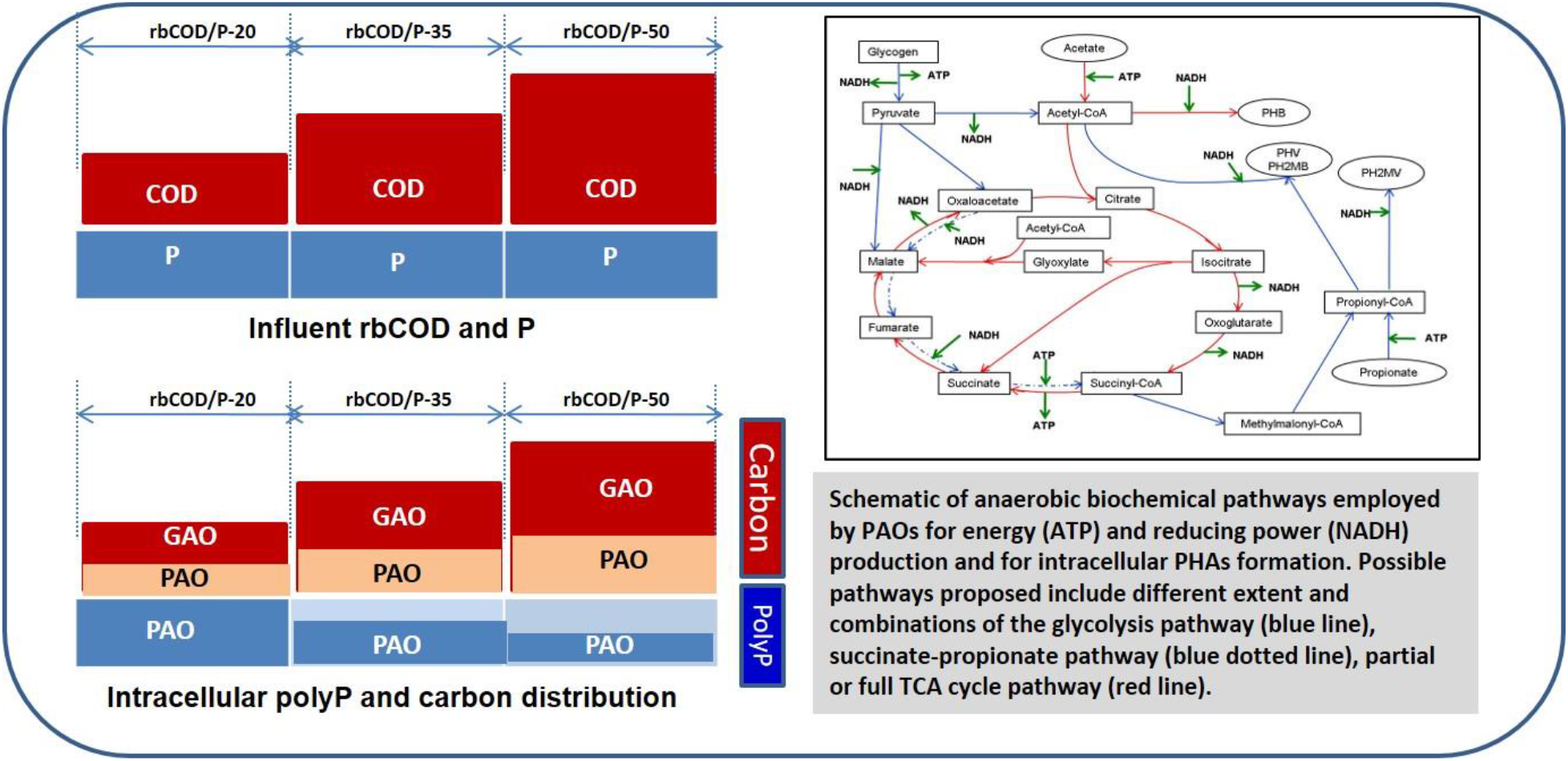

## 1. Introduction

The increasingly stringent limits imposed on wastewater effluent phosphorus demand for higher-level treatments through more reliable and better optimization of phosphorus removal processes. Although enhanced biological phosphorus removal (EBPR) process is considered a potentially efficient process with economic and environmental advantages compared to traditional chemical phosphorus removal, the benefits are often offset, in practice, by the needs to have standby chemicals for achieving reliable and consistent performance. There is still knowledge gap in understanding the mechanism and factors that control the stability of the process, particularly for achieving extremely low effluent limits and, as a result, process stability and system performance have been seen to vary among facilities (Gu et al., 2005; Neethling et al., 2005; Stephens et al., 2004).

EBPR performance and stability have been shown to be affected by many factors among which, the competition of the two main functionally relevant populations, namely polyphosphate accumulating organisms or PAOs and glycogen accumulating organisms or GAOs, were found to be crucial for achieving successful operation of EBPR. Although deterioration of EBPR performance has been attributed to the proliferation of GAOs in lab-scale EBPRs and also at full scale EBPR facilities (Gu et al., 2005; Cech and Hartman, 1993; Saunders et al., 2003), others (Gu et al., 2008; Tu and Schuler, 2013) have also shown that efficient EBPR could also be achieved with relatively high abundance of GAOs being present, highlighting the need for further understanding of the impact of GAOs on EBPR. In addition, the phylogenetic identities, phenotypic elasticities, and biochemical metabolic pathways involved in these two groups of microorganisms are still not fully understood. There are still uncertainties regarding the mechanism and metabolic pathways in the EBPR process; particularly, the involvement of either TCA cycle or glycolysis pathway (Entner-Doudoroff versus Embden-Meyerhof-Parnas pathway) for energy and reducing power generation and, the extent of possession and utilization of these pathways by different phylogenetic PAOs or GAOs groups under various conditions remain unclear (Zhou et al., 2010). Interestingly, recent studies pointed out that these two groups could change and switch their phenotypes and metabolic pathways under different conditions revealing their metabolic flexibility (Acevedo et al., 2012; Lanham et al., 2013; daSilva et al., 2018).

Understanding and designing the conditions that are favorable for PAOs over GAOs and other competing microorganisms is considered necessary to maintain the system stability and performance (Neethling et al., 2005; Barnard et al., 2005). Several factors have been shown to impact the competition between PAOs and GAOs, including influent bio-available carbon to P ratio, solid retention time, VFA loading rate, feeding strategy and composition, hydraulic retention time, temperature, pH, dissolved oxygen and, salinity etc. (Cech at Hartman, 1993; Tu and Schuler, 2012; Whang and Park, 2006; Liu et al., 1997; Filipe et al., 2001; Lopez-Vasquez et al., 2009). Among these, the influent C/P ratio is particularly important since it has been correlated with EBRP performance and stability (Gu et al., 2008). The observed stoichiometric requirement of carbon for a unit amount of phosphorus to be removed has been around 10-20 mg rbCOD/mg P removed (Barnard et al., 2005; Tchobanoglous, 2003). Higher rbCOD/P ratios (40-50 mg-rbCOD/mg-P) have been seen to be associated with GAO dominated culture (Liu et al., 1997; Broughton et al., 2008; Kong et al., 2002; Schuler and Jenkins, 2003; Oehmen et al., 2007) and lower ratios (<10-20 mg-rbCOD/mg-P) have been associated with PAO dominated culture. The range of rbCOD/P ratio for satisfying P removal in Water reclamation facilities was recommended as 15:1-25:1 (Randall et al., 1992; Tetreault et al., 1986). Gu et al (2008) hypothesized that although the excessive amount of available carbon can harbor the proliferation of GAOs, stable process can be maintained as long as the operational conditions are controlled to kinetically favor the growth of PAOs over GAOs.

The link of influent rbCOD/P ratio with microbial population structures and consequent impact on EBPR performance stability warrants further investigation. Specifically, the impact of varying rbCODd/P loading conditions on the relative abundances of both PAOs and GAOs along with their metabolic changes and competition requires better understanding. The unavailability of diverse PAO and GAO isolates, and the lack of tools to monitor the metabolic state of these two key populations in a mixed culture, make it often difficult to understand the influence of process parameters on the specific population levels. A single cell Raman micro-spectroscopy method that allows for phenotype-based quantification of relative PAO and GAO abundance, simultaneous intracellular detection and quantification of polymers (i.e. polyphosphate, PHB and glycogen) at individual cellular level was developed by Majed et al. (2009, 2010). Later on, Raman-based phenotypic operational phenotype unit (OPUs) was also linked with operational taxonomic unit (OTUs) by Li et al (2018). In this study, we further employed the developed Raman microscopy method to evaluate the impact of influent rbCOD/P ratio on the EBPR system, including overall performance, relative PAO/GAO population abundance and, particularly, on the intracellular polymer dynamics at both single cellular and population levels. These results provided new insights into metabolic diversity, involvement of biochemical pathways and the mechanisms of EBPR.

## 2. Material and Methods

### 2.1 Sequencing batch EBPR reactors

Three identical lab-scale SBR-EBPR systems were operated with three different influent rbCOD/P ratios of 20, 35 and 50 (ratios relevant to full-scale EBPR processes in US), respectively by changing the influent COD concentrations in relative to constant influent orthoP level of 10 mg-P/L. The SBRs were controlled at a constant room temperature of 19-22°C and operated with four six-hour cycles per day with each cycle consisting of: 10 minutes’ fill followed by 130 minutes of anaerobic phase, 183 minutes of aerobic phase, 30 minutes of settling and then 7 minutes of withdrawing. The composition of synthetic wastewater feed was according to Schuler and Jenkins (2003). Phosphorus was added as 45 mg/L sodium phosphate monobasic (NaH_2_PO_4_ • H_2_O) (10 mg-P/L). Influent organic feeding ranging from 200 – 500 mg COD/L as sodium acetate (CH_3_COONa • 3H_2_O) was provided with supplement of 15 mg/L of casamino acids. We recognize that this study only focused on acetate-fed system that are dominated by *Accmulibacter*-like PAOs and investigation of EBPR systems with more complex carbon feed (i.e mixture of acetate and propionate) is warranted for future studies. Nitrogen was added as ammonium chloride (NH_4_Cl) to maintain stoichiometric requirement of nitrogen for growth (rbCOD:N:P of 100:5:1). Allylthiourea was added at 2 mg per liter of feeding to inhibit nitrification. The SRT and HRT of the system were maintained at 7 days and 12 hours respectively. For each rbCOD/P ratio, the SBR was operated for duration of at least 3 times SRT (21-25 days) for obtaining stable performance before subjecting to populations’ study and Raman analysis.

### 2.2 EBPR performance evaluation

Three SBRs were operated with rbCOD/P ratios of 20, 35 and 50 respectively. For each rbCOD/P ratio operations, P release and uptake cycles were monitored during the steady state period to determine the EBRP activities, including P release and uptake rates, P release to acetate uptake ratios and glycogen degradation to acetate uptake ratios. Samples (in duplicate) were taken at the beginning and at the end of anaerobic cycle and at the end of aerobic cycle for each of the three SBRs with varying influent rbCOD/P ratios for Raman analysis, as well as for quantifying cellular-level intracellular polymers content including polyphosphate, PHB and glycogen. In addition, samples were collected at 15-60 minutes’ intervals for bulk chemical analysis to be performed for soluble orthophosphate, total phosphate, acetate, total solids and glycogen content in sludge. The filtered samples through 0.45 μm filter papers were analyzed for orthophosphate (orthoP; (PO_4_^3^)^−^) and acetate (CH_3_COO^−^) using DX-120 ion chromatograph (Dionex Benelux, Belgium). All phosphorus fractions were measured according to the standard method (4500-P) (APHA, 1998). Glycogen was measured according to the method specified by Erdal (2002).

### 2.3 PAOs/GAOs population analysis

Presence of PAOs in the reactor was confirmed with phosphate removal performance evaluation, Neisser and DAPI staining (Jenkins et al., 1993; Streichan et al, 1990) for total PAO observation and, FISH for detecting known candidate PAOs and GAOs. Oligonucleotide probes targeting *Accumulibacter* PAOs, *Actinobacteria* PAOs, *Competibacter* GAOs, *Defluvicoccus* clusters 2 GAOs were used in FISH analysis (Detailed listing of probes is provided in STable 1). The staining, FISH protocol and hybridization conditions used were previously described (He et al., 2008; Zilles et al., 2002). DAPI staining for polyP was carried out at 50 μg/ml of DAPI for 1 min, whereas, DAPI staining for total population in hybridized slides (to analyze for relative abundance of phylogenetic sub-groups of PAOs and GAOs over total population) was carried out at 1μg/ml of DAPI for 10 min. Hybridized and DAPI stained cells were observed with an epifluorescent microscope (Zeiss Axioplan 2, Zeiss, Oberkochen, Germany). For the quantification of the relative proportion of the target types of cells, around 20 micrographs were collected with random fields of view from the same slide/sample and average abundance of the target cells was calculated as the relative proportion of fluorescing area having the target label (PAO/GAO) compared to the area of the total population (DAPI) using DAIME (Digital Image Analysis in Microbial Ecology) software version 1.3.1 (http://www.microbial-ecology.net/) (Daims et al., 2006). The standard error of the mean (SEM) was calculated as the standard deviation of the area percentages divided by the square root of the number of images analyzed. For Raman analysis, fractions of target populations (PAOs/GAOs) were determined by averaging the fractions of samples from aerobic phase according to Majed et al (2012).

### 2.4 Raman Micro-Spectroscopy Analysis

Samples subjected to Raman analysis were prepared on optically polished CaF_2_ windows (Laser Optex, Beijing, China) according to Majed et al (2010). Samples were homogenized rigorously by 26-gauge needle and syringe to obtain a more uniform sample. Raman spectra for at least 40-45 single cells were examined for each sample and the sample size was determined with consideration of both the desired accuracy and labor-intensiveness of the Raman analysis. Statistical analysis of the sample size and validation of the analysis accuracy and reliability was demonstrated in our previous publications (Majed et al., 2010; Li et al., 2018) and also provided in supporting information (SFigure 3 and SFigure 4). Raman spectra were acquired using a WITec, Inc. (Ulm, Germany) Model CRM 2000 confocal Raman microscope. Excitation (ca. 30 mW at 633nm) was provided by a Helium/Neon laser (Melles Griot, Carlsbad, CA). Details on the acquisition of spectra can be obtained in Majed et al. (2009). Relative quantity of polyP content in each individual cell was evaluated based on the Raman intensity (peak height in the unit of Charged Coupled Device (CCD counts) of the PO_2_^−^ stretching band occurring around 1168-1175 cm^−1^ wave number region after background correction (Majed et al., 2009). The C=O stretching band of ester linkage occurring around 1734 cm^−1^ and glycogen vibration occurring around 480 cm^−1^ were used for quantification of PHB and glycogen content, respectively (Majed and Gu, 2010). Total PHV concentration associated with the two populations could not be quantified via Raman due to the overlapping of peak positions between PHV and glycogen (Majed and Gu, 2010). Depending on the time point during the anaerobic/aerobic EBPR cycle when the sample was taken, and the expected corresponding Raman polymers spectrum based on current understanding of the EBPR mechanisms and polymer functions, cells containing polyP, with or without other polymers (glycogen, PHA) were categorized as PAOs and the cells containing only glycogen or combination of glycogen and PHA, were assigned as GAOs. Detailed description of the rationale and validation of the proposed quantification methods is referred to Majed et al. (2012).

## 3. Results and Discussion

### 3.1 Impact of Influent rbCOD/P on EBPR performance and kinetics

Table 1 shows the EBPR activities-related stoichiometry observed in EBPR systems along with the performance and stability of the system at different influent feeding rbCOD/P ratios. Operational data for the SBRs under monitoring (STable 2) indicated that percent removal of P changed from 82% to 95% to 97% as rbCOD/P ratio changes from 50 to 35, then to 20 respectively. The results showed good P removal at rbCOD/P ratio of 20 to 35, and significant decrease in P removal efficiency at rbCOD to P ratio of 50.

**Table 1:** P removal performance and activity data of the SBR-EBPR systems operated with influent rbCOD/P ratio at 20, 35 and 50, respectively.

| Influent<br>mg-COD/m<br>g-P | Cumulative<br>Frequency<br>Effluent<br>ortho-P<<br>1mg P/L | Cumulative<br>Frequency<br>of Effluent<br>ortho-P<<br>0.5mg P/L | P<br>Removal<br>Efficiency<br>(%) | P release<br>rate, mg-P/gVSS.h | P uptake<br>rate,<br>mg-P/gVSS.<br>h | P<sub>release</sub>/HAc<br>uptake<br>Pmol/Cmol | Gly<sub>utilization</sub>/<br>HAc<sub>uptake</sub><br>Cmol/Cmol | P content,<br>mg-P/mg-VSS |
| --- | --- | --- | --- | --- | --- | --- | --- | --- |
| **20** | 0.85 | 0.6 | 97 | 50.63 | 23.81 | 0.6 | 0.41 | 0.09 |
| **35** | 0.86 | 0.63 | 95 | 38.14 | 21.2 | 0.47 | 0.74 | 0.06 |
| **50** | 0.5 | 0.31 | 82 | 4.38 | 9.82 | 0.22 | 1.04 | 0.024 |

Through comparison of cumulative frequency of the occurrence that effluent orthoP is obtained below <1 mg/L and below <0.5 mg/L, the system with COD/P ratio of 35 ensures the maximum stability while the system with rbCOD/P ratio of 20 performs almost similar. However, system is far from stable for the SBR system with rbCOD/P ratio of 50.

According to established EBPR models, EBPR metabolic pathways demand certain stoichiometric relationships among the storage polymers. The P_relaease_/HAc_uptake_ (P/HAc) ratio is often used as an indicator of the relative PAO and GAO activities and abundance. As shown in Table 3, the P/HAc ratio was 0.6 Pmol/Cmol for rbCOD/P ratio of 20 which decreases linearly with increasing rbCOD/P ratio. Previous studies indicated that higher influent rbCOD/P led to more GAO-dominant microbial population in the system (Smolders et al., 1994). This is confirmed and supported by our population analysis results that are discussed in the section 3.2.

The anaerobic glycogen_utilization_/HAc_uptake_ ratios (Gly/HAc ratio) decreased linearly from 0.41 to 1.04 Cmol/Cmol from rbCOD/P ratio of 20 to 50 respectively. These ratios are within the range between TCA cycle and glycolysis activity, as reported in full-scale EBPR facilities and also in established models for PAOs and GAOs (Gu et al., 2008; Lanham et al., 2013; Smolders et al., 1994; Zeng et al., 2003). The increase of Gly/HAc indicates a preferential reliance by the population on TCA cycle at a lower rbCOD/P to an increased reliance on the glycolysis pathway at a higher rbCOD/P. Similar scenario has been observed in several EBPR facilities in Denmark (Lanham et al., 2013). In addition, decrease in the overall P content from 0.09 to 0.024 mg-P/mg-VSS (with increasing rbCOD/P) indicate a decrease in the relative population of PAOs with increasing rbCOD/P ratio.

### 3.2 Impact of influent rbCOD/P on relative PAO- GAO population abundance and Association with EBPR activities

Figure 1 shows the PAOs (Figure 1a) and GAOs (Figure 1b) population fractions changes in response to varying influent rbCOD/P ratios, using conventional DAPI staining, FISH measurements and Raman analysis. As shown in Figure 1a, total PAOs determined via Raman spectrum agreed well with those estimated by DAPI staining for all rbCOD/P ratios. Currently, there is no other method available for quantifying total GAOs in EBPR systems; therefore, the GAOs abundance determined by Raman could not be compared. Note that the sum of GAOs and PAOs, as quantified by Raman analysis, comprised around 85-90% of total microbial population in the SBR-EBPR studied, indicating that our methods captured majority of the population.

**Figure 1:**
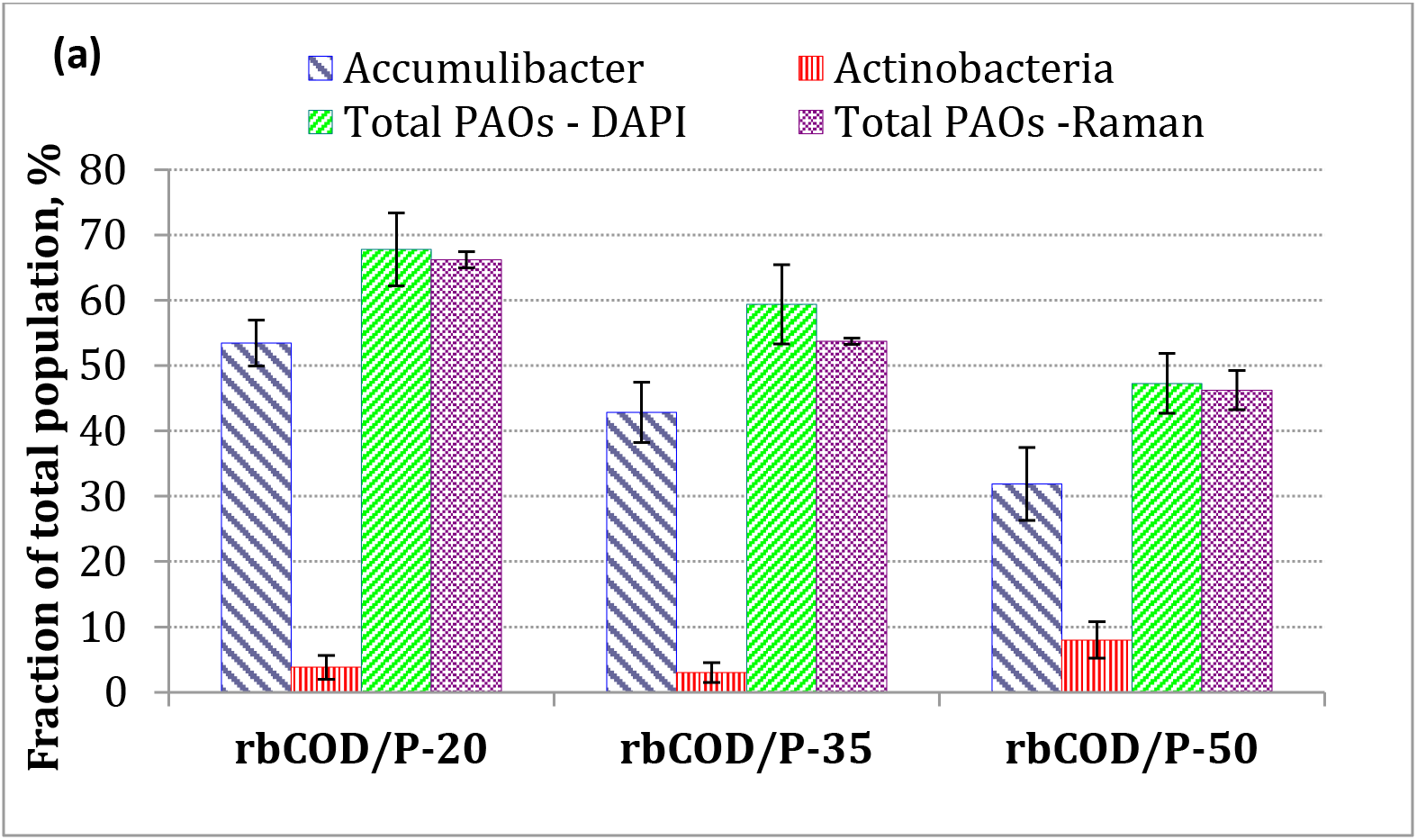
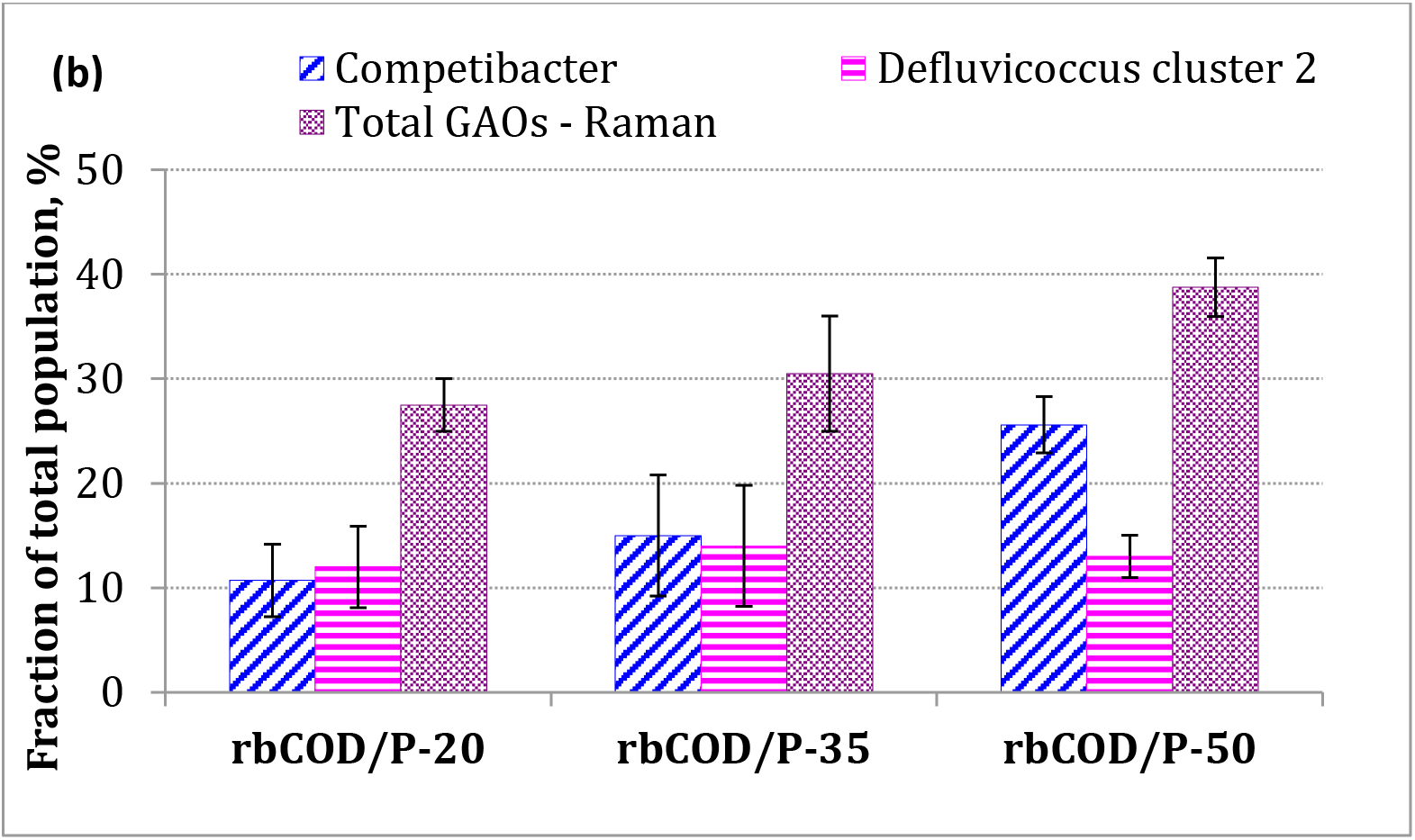
Abundances of population fractions belonging to the groups (a) PAOs and (b) GAOs at different rbCOD/P ratios quantified via DAPI staining (total PAOs), FISH (known candidate sub-PAOs and GAOs) and Raman polymers spectrum analysis (total PAOs and GAOs based on intracellular polymers). Error bars represent standard error

Both *Accumulibacter*-like PAOs and total PAOs abundance decreased, as the influent rbCOD/P (mg-COD/mg-P) ratio increased from 20 to 50 (Figure 1a). *Accumulibacter* PAOs comprised around 70% of total PAOs population at all rbCOD/P ratios studied. *Actinobacteria* type PAOs constituted a minor fraction of total PAOs being less than 5% of the total population at rbCOD/P ratios of 20 and 35, however, their abundance increased to 8% at rbCOD/P of 50. *Actinobacteria* PAOs are known to grow on amino acids instead of acetate and the casamino acids in the feeding likely provided the amino acid source that supported the growth of *Actinobacteria*. Concurrently with the decrease of relative PAOs abundance, the relative GAOs abundance increased by 40% with the increasing rbCOD/P ratios from 20 to 50 as shown in figure 1b. *Alphaproteobacterial Defluvicoccus* cluster 2 (*DF2*) and *Competibacter* type GAOs accounted for majority (>80%) of the total GAOs (based on Raman measurements since there is currently no other method available for quantifying total GAOs) in our culture (Figure 1b). *Competibacter* abundance increased by >40% with the increasing rbCOD/P ratios from 20 to 50, but abundance of DF2 did not seem to vary, indicating that *Competibacter* was likely the one that responded to the change in loading ratio. Muszyński and Miłobędzka (2015) also observed increase in *Competibacter* abundance from 4 to 20% when the rbCOD/P ratio was changed from 15:1 to 100:1 with aerobic granular sludge while abundance of *Alphaproteobacterial Defluvicoccus* cluster 1 (*DF1*) remained constant at 2% (they did not detect DF2 type GAOs in the granular sludge). These results clearly showed that both relative abundance and community compositions of PAOs and GAOs shifted as the influent rbCOD/P ratio changed.

The changes in relative abundance of PAOs and GAOs in response to influent rbCOD/P variations were consistent with the chemical analysis of P content level in the sludge and the evaluation of EBPR activities, as summarized in Table 1 (profiles demonstrated in SFigure 1). With a higher carbon loading with respect to phosphorus, the P content in sludge, the P release rate and the P_release_/HAc_uptake_ ratios decreased, indicating a decline of relative PAOs activities (presumably proportional to PAO populations) and increase in relative GAOs in the sludge. Correlation analysis of relative PAOs abundance (%) with the EBPR activities parameters seemed to indicate that relative total PAOs abundance change correlated more significantly(p<0.05) with P_release_/HAc_uptake_ ratio (r = 0.99, p = 0.0351) and sludge P content (r = 0.99, p = 0.0468), but less so with P release rate (r = 0.99, p = 0.0991), glycogen_degradation_/HAc_uptake_ ratio (r = −0.99, p = 0.0836), and P uptake rate (r = 0.97, p = 0.155). The result also showed positive correlation between the P_rel_/HAc_uptake_ ratio and the abundance ratio of *Accumulibacter*/*(Competibacter*+DF2) (correlation coefficient, r = 0.97; p = 0.1553), as well as P_rel_/HAc_uptake_ ratio and total PAO/GAO (analyzed by Raman) (r = 0.98, p = 0.1248) at different rbCOD/P ratios.

Previous studies have indicated that rbCOD/P ratio seem to dictate the relative PAO and GAO population abundance based on indirect observation of changes in EBPR activities (since total GAOs could not be quantified previously) (Gu et al., 2008; Liu et al., 1997; Schuler et al., 2003). Our results are consistent with the previous studies, and more clearly demonstrated the impact of rbCOD/ P ratio on both total and different sub-populations of PAO and GAOs, as well as their associated phenotypic activities. We recognize that this study only focused on acetate-fed system that are dominated by *Accmulibacter*-like PAOs and investigation of EBPR systems with more complex carbon feed (i.e mixture of acetate and propionate) is warranted for future studies.

One intriguing observation is that P removal performance deteriorated at rbCOD/P ratio of 50, even though the system still contained rather high relative abundance of total PAO and *Accumulibacter*-like PAOs (47%, 32% respectively.), which are higher than typical range observed at full-scale EBPR systems (Neethling et al., 2005; Gu et al., 2008; Lanham et al., 2013; Gu et al., 2018). This suggests, in agreement with previous observations, that EBPR performance does not always correlate with relative PAOs abundance alone.

### 3.3 Impact of Influent rbCOD/P ratio on population-level distributions of intracellular polymer content in PAOs and GAOs

Raman analysis allows for the quantification of intracellular inclusion of polyP, PHB and glycogen within each individual cell and therefore can reveal the level and distribution of intracellular polymer contents among populations. In addition, by identifying the cells as either PAOs or GAOs based on their unique polymer combination as described previously, the changes and dynamics of the functionally relevant intracellular polymers associated with either PAOs or GAOs can be, for the first time, evaluated separately. However, it should be noted that evaluation of the levels of intracellular polymeric inclusions and the resulting distributions do not necessarily correspond to the rate of increase or decrease of polymer formation or degradation at any given condition, nevertheless the variations in the levels at different conditions depict the dynamics of intracellular transformations in response to the influent loading changes.

Figure 2 and Figure 3 show the comparison of quantity level and distributions of different intracellular polymers in PAOs for EBPR systems operated under three different influent rbCOD/P ratios. Both 50 percentile and maximum polyP intensity inside individual PAO cells decreased as the rbCOD/P ratio increased even though the influent P concentration remained the same (SFigure 1). Drastic decrease of cellular level polyP content at increased rbCOD/P ratio is quite contrary to the traditional assumption in EBPR models that considers that the maximum saturated intracellular polyP levels is relatively constant and only PAO populations abundance varies with the external condition (Streichan et al., 1990). Therefore, PAOs abundance changes based on the bulk sludge P content or overall P release during EBPR process may not be adequate. In contrast to significant polyP abundance changes, the intracellular PHB levels in the PAOs exhibited slight increase (38% for median, 37% for maximum values disregarding the outlier) as the influent rbCOD/P ratio increases from 20 to 50 (Figures 2 and 3). There was no significant change in the median level of intracellular glycogen level in PAOs, however maximum and minimum levels increased (25% and 43%respectively, Figure 3) as rbCOD/P increased from 20 to 50. This warranted the one-way ANOVA test among the polymeric inclusion data which revealed F_polyP_(2,301) = 53.6, p≪0.05; F_PHB_(2,92) = 6.87, p≪0.05; F_glycogen_(2, 119) = 10.5, p≪0.05 for PAOs suggesting that the changes in polymeric inclusion levels with changes in rbCOD/P ratio are statistically significant. This implies that the intracellular polymers content and their stoichiometric ratios in PAOs are rather dynamic depending on the influent carbon loadings, which can be associated with both PAO phylogenetic diversity and phenotypic elasticity. Therefore, the saturation level (maximum) of these intracellular polymers that are considered to be a constant in current EBPR models (Schuler, 2005) needs further justification.

**Figure 2:**
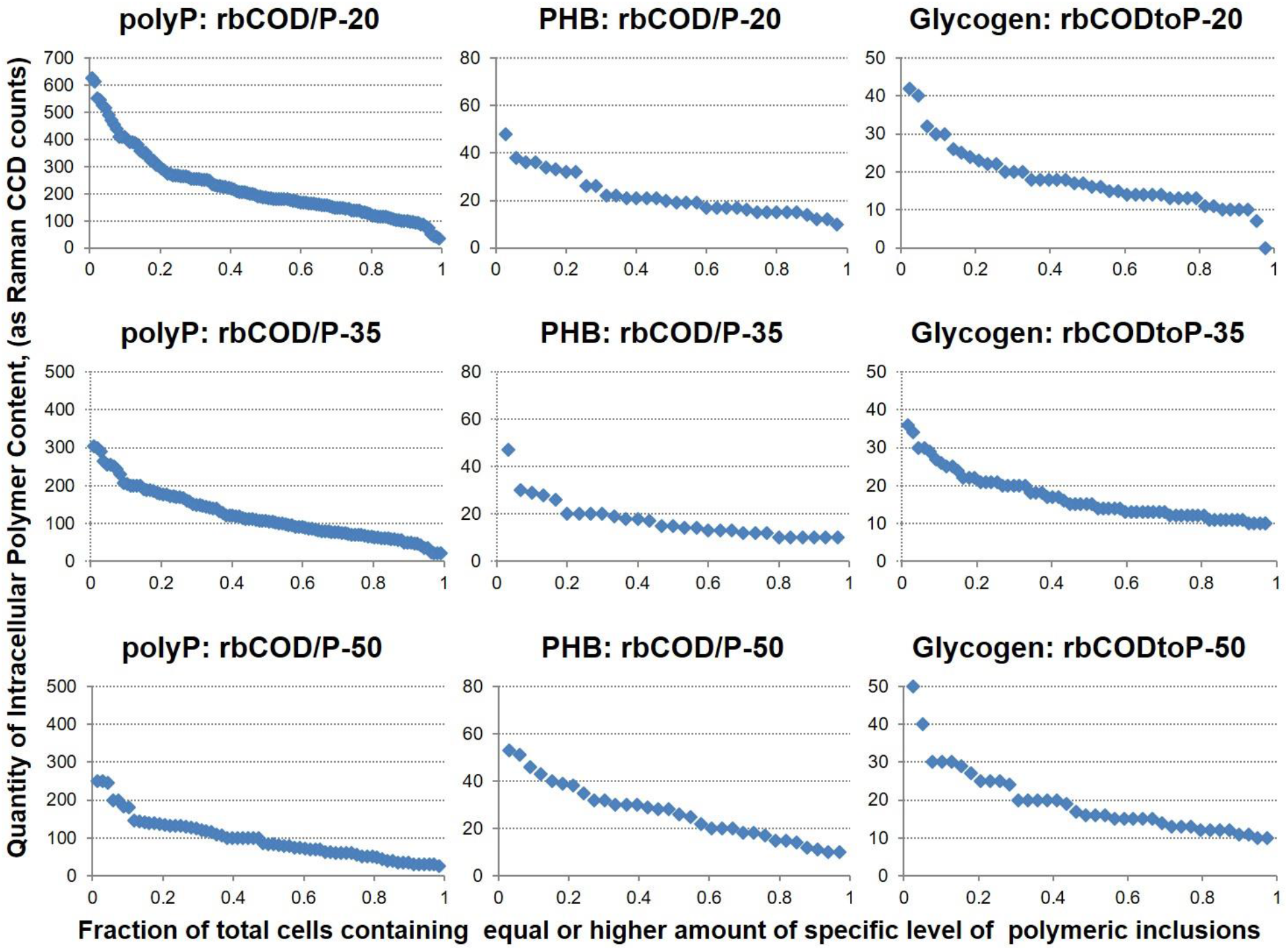
Intracellular polymer content level (intensity as CCD counts) distribution among PAOs cells for polyphosphate, PHB, and glycogen quantities, respectively. Data based on batch testing sampling of all cells subjected to single cell Raman micro-spectroscopy analysis in the three EBPR SBR systems fed with different influent COD/P (mg/mg) ratios. *X* axis: fractions of cells that contained equal or higher amount of the specific level of polymeric inclusion at any given testing. Y-axis: level of polymeric inclusion in CCD counts.

**Figure 3:**
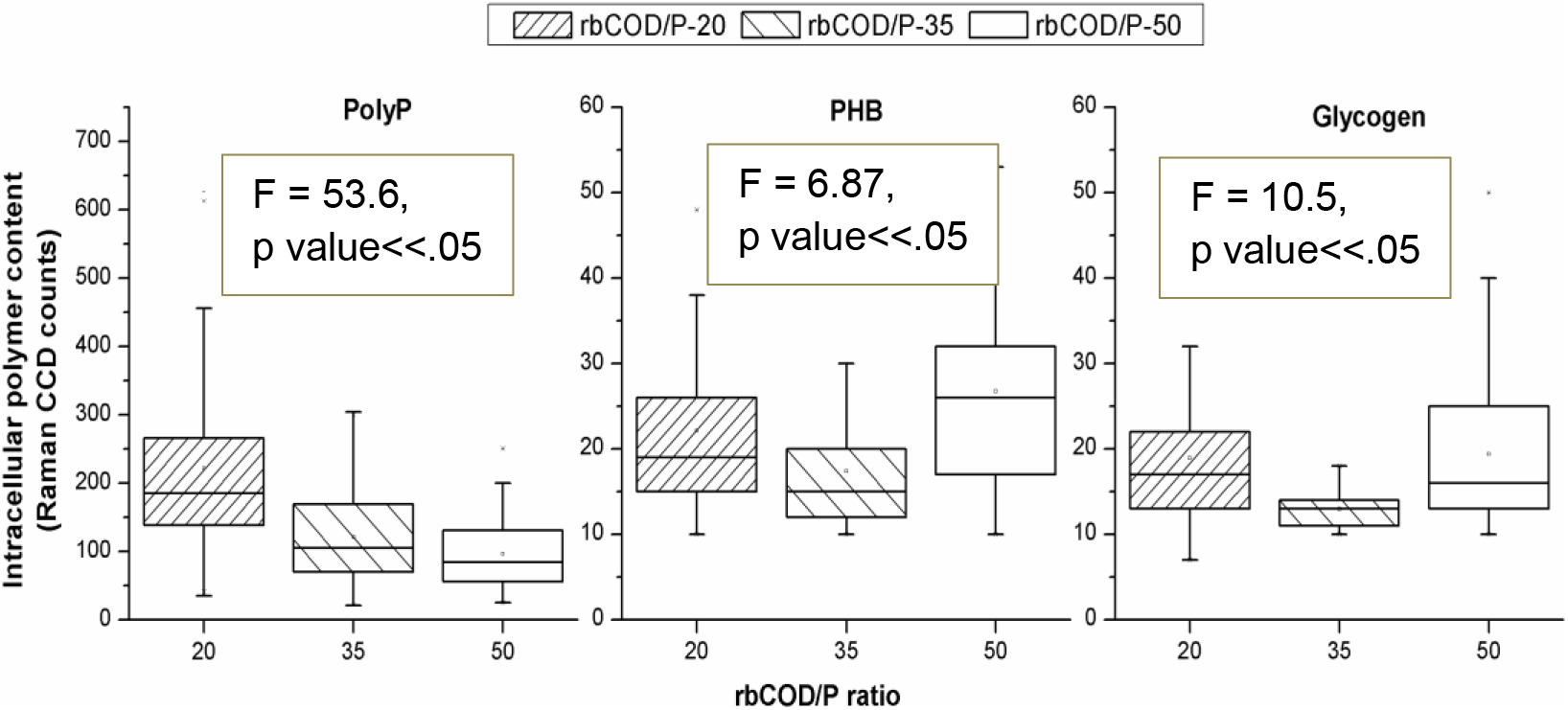
Box plots showing the minimum, maximum and median levels of intracellular polymer content levels of polyP, PHB and glycogen polymers in individual PAO cells revealed by single cell Raman micro-spectroscopy (respective F statistic and P values are shown in figures)

Similarly, observations of inclusion level and distribution for intracellular glycogen and PHB content in individual glycogen containing cells (GAOs) were revealed by single cell Raman microspectroscopy (Figure 4 and SFigure 2). Significant increase in cellular PHB content (50% increase in median and 31% increase in maximum cellular level) with elevation of rbCOD/P ratios (from 20 to 50) in GAO cells (Figure 4) was observed. Intracellular glycogen level in GAOs exhibited overall increase for maximum levels (50% increase from rbCOD/P of 20 to 50), however median values remained almost similar. This warranted the one-way ANOVA test among the polymeric inclusion data which revealed F_PHB_ (2,96) = 5.5, p≪0.05; F_glycogen_ (2,171) = 1.72, p>0.05 for GAOs. This suggests the higher carbon loading to the EBRP systems not only resulted in increase of relative abundance of GAOs, but also led to either selection of GAOs with higher intracellular PHB, and/or encouraged GAOs to accumulate more PHA pool inside cells. However, the increment of glycogen pool within GAO cells with increase in rbCOD/P ratio is not statistically significant.

**Figure 4:**
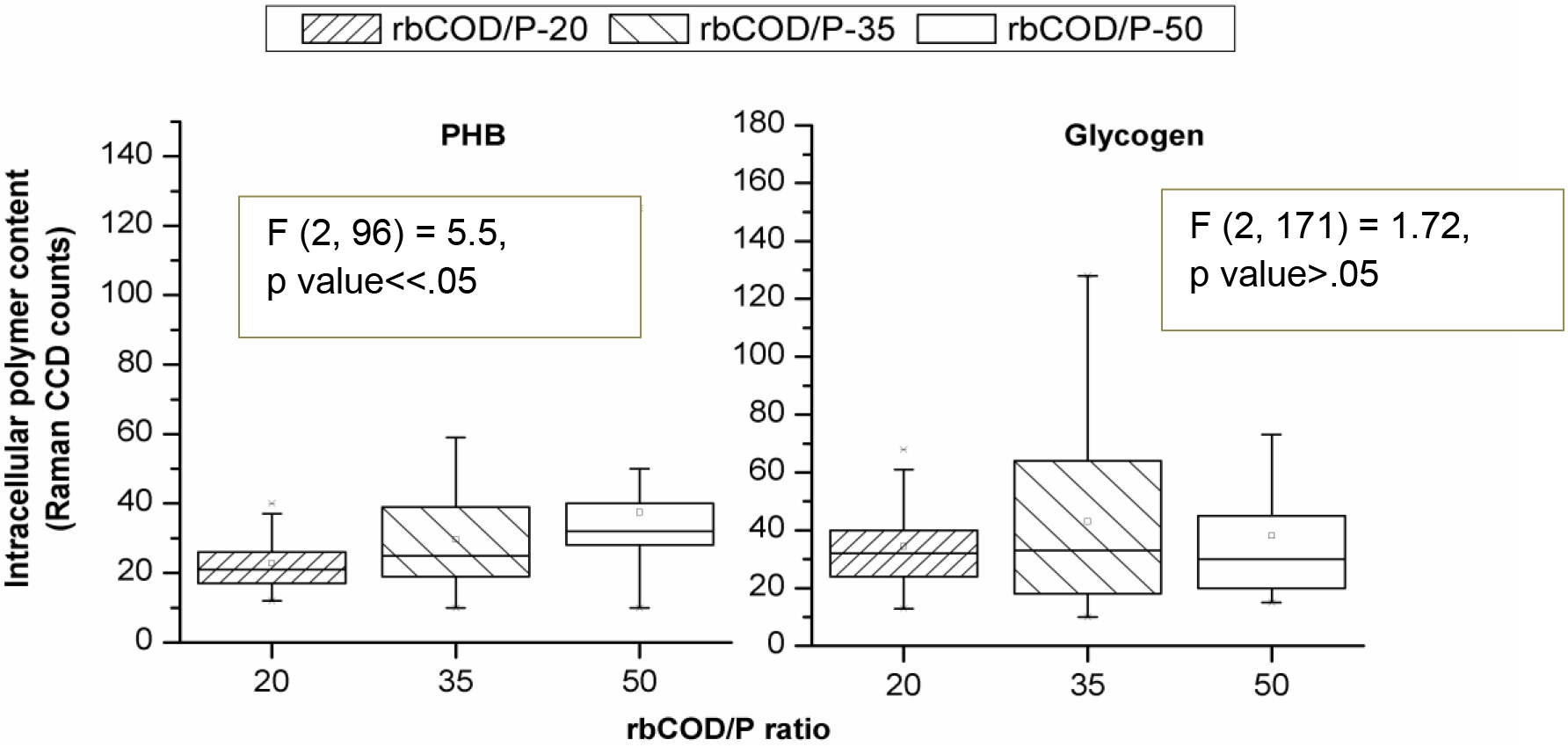
Box plots showing the minimum, maximum and median levels of intracellular polymer content levels of PHB and glycogen polymers in individual GAO cells revealed by single cell Raman micro-spectroscopy (respective F statistic and P values are also mentioned)

These results, for the first time, revealed the individual cellular level polymers level changes in both PAOs and GAOs populations in response to changes in influent carbon and P loading conditions. These observed intracellular polymers dynamics could result from and reflect the changes in phylogenetic diversity and, or metabolic functions shifts in PAOs, which requires further investigation. The results also imply that there is great heterogeneity of the intracellular polymers storage amount among PAOs and GAOs, which can be dynamic depending on operational conditions.

### 3.4 Distribution of Carbon and polyP between PAOs and GAOs at varying COD/P condition

Using the Raman-based PAO and GAO identification methods described earlier, differentiated intracellular polymer content associated within PAOs versus those within GAOs could be quantified and evaluated separately (Majed and Gu, 2010). Figure 5a shows the total and distributed PHB (normalized amount at the end of the anaerobic phase when the PHB content is presumably to be at the maximum level) and glycogen content levels (normalized amount at the end of the aerobic phase when the glycogen content is presumably to be at the maximum level) inside PAOs and GAOs. Figure 5b shows the ratio of PHB and glycogen content in PAOs in relative to those in GAOs at different rbCOD/P loading conditions and figure 5c is a qualitative illustration of the distribution and carbon flow among PAOs and GAOs in response to the increase in influent carbon loads.

**Figure 5:**
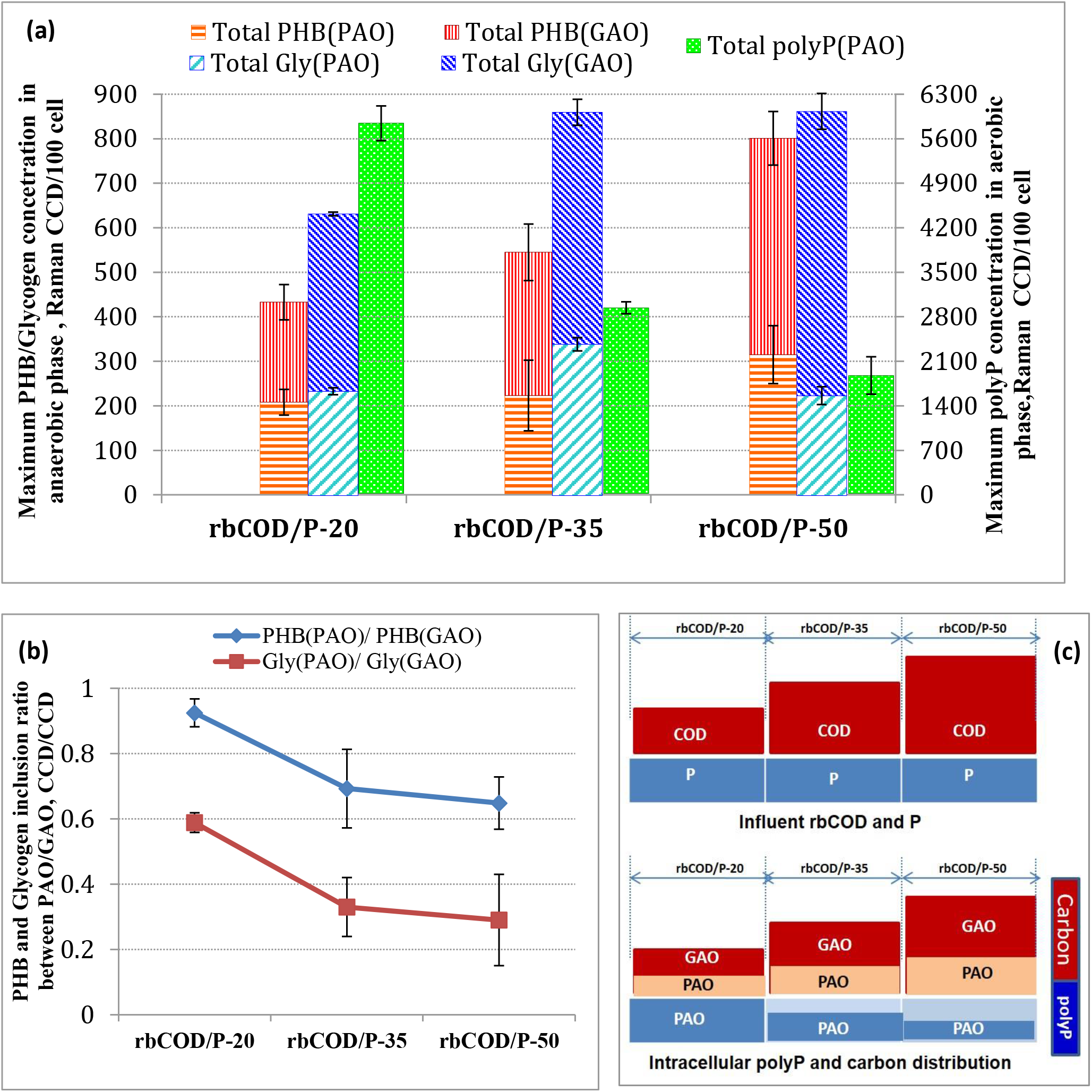
(a) Total and distributed intracellular PHB, glycogen and polyP content between PAOs and GAOs in EBPR systems operated with different influent rbCOD/P ratios. The concentration of the polymer inclusions in a given biomass sample was determined as the sum of total polymers amounts in those cells who contain the polymer(s) normalized by the number of total cells analyzed (sample size). PHB, glycogen and polyP content in a biomass sample was measured with samples taken from anaerobic, aerobic and aerobic phases respectively; (b) Ratio of the PHB and glycogen within PAOs to those in GAOs at different COD/P ratios. (Error bars represent the standard deviation between two separate measurements); (c) Qualitative illustration of intracellular carbon storage distribution between PAOs and GAOs in response to varying carbon loading conditions.

Both total PHB and total glycogen concentration in the overall mixed population (sum of those in both PAOs and GAOs) exhibited elevating trends with the increasing rbCOD/P ratio. Majority portion of incremental PHB and glycogen content was associated with GAOs, indicating the flow of the increased carbon to GAOs as a result of increased relative abundance of GAOs as well as elevated average individual cellular glycogen content in GAO cells. The ratio of PHB inside PAOs to that inside GAOs dropped from 0.92 to 0.65 when rbCOD/P ratio increased from 20 to 50, (Figure 5b). These results clearly demonstrated the quantitative shift in intracellular carbon storage distribution between the two populations, from having a good portion (50%) of the total PHB in PAO populations to having majority of the carbon shuttled to GAOs as the rbCOD/P increased. These results support the hypothesis and carbon distribution model proposed by Gu et al (2008) that as the influent rbCOD/P ratio increases, the excessive carbon beyond stoichiometric requirement for PAOs would be diverted into GAOs.

Also shown in Figure 5a is the normalized polyP concentration (total polyP amount determined by Raman normalized to sample size) that exhibited dramatic decrease with increase in rbCOD to P ratio as a result of decrease in individual cellular polyP intensity. These results are consistent with bulk P content (Table 1), suggesting again the validity of the Raman analysis for polyP. The quantitative measurement of intracellular polymers associated with PAO and GAO populations allowed for the estimation of changes in the stoichiometry of the utilization and formation of these polymers with different loading conditions. Table 2 summarizes the ratios of PHB formation to polyP utilization, glycogen utilization to polyP utilization and PHB formation to glycogen utilization during the anaerobic phases in PAOs. As described in our previous studies, the lack of standard chemicals for polymers and difficulty with Raman method to establish a reference calibration for a mixed culture (activated sludge) as ours did not yet allow the conversion of CCD count to conventional units of mg/L (Majed et al., 2009). Therefore, we could not compare our stoichiometric values directly with those reported in the literature yet (Oehmen et al., 2010). Nevertheless; the ratios used here are still valid for assessing the impact of rbCOD/P ratio on these values. Based on the current biochemical model of EBPR, the theoretical stoichiometric ratios between polyP, PHB and glycogen polymers are generally assumed to be constant for a given pathway within a given population (Smolders et al., 1994). However, as shown in Table 2, each of the ratios varied consistently for PAOs as the loading conditions changed. These results, for the first time, demonstrated that the EBPR stoichiometry could vary significantly with loading conditions, possibly due to the metabolic states changes (e.g utilization of different pathways) and /or population changes within phylogenetic sub-groups (e.g sub clusters of *Accumulibacter*-like PAOs).

**Table 2:** Stoichiometric ratios of polyP utilization, PHB formation and glycogen utilization in the anaerobic phase for PAOs, in SBR-EBPR system operated with different influent rbCOD/P (mg/mg) ratios.

|  | PHB formation /<br>polyP utilization | Glycogen utilization<br>/polyP utilization | PHB formation<br>/Glycogen utilization |
| --- | --- | --- | --- |
|  | CCD/CCD | CCD/CCD | CCD/CCD |
| **rbCOD/P-20** | 0.044 | 0.009 | 4.84 |
| **rbCOD/P-35** | 0.13 | 0.07 | 1.81 |
| **rbCOD/P-50** | 0.24 | 0.21 | 1.21 |

### 3.5 Insights into the metabolic pathways involved in EBPR

Uncertainties still exist regarding the pathways employed by PAOs for reducing power generation in anaerobic EBPR process, which would affect the stoichiometric ratios and the amount and type of PHA formation from acetate uptake (Zhou et al., 2010). Current understanding indicates that PAOs can use glycolysis in addition to partial TCA cycle (reductive branch of TCA cycle, also called succinate-propionyl-coA pathway), split (reverse) TCA cycle and/or glyoxylate shunt based on carbon preservation under different polymer content (daSilva et al., 2018; Martin et al., 2006; Wilmes et al, 2008; He et al., 2010; Zhou et al., 2009) (provided as SFigure 5 and SFIgure 6). The extent of involvement and utilization of these pathways also vary depending on the phylogenetic subgroups of PAOs and different external loading/nutrient availability conditions (Zhou et al., 2010; daSilva et al., 2018; Oehmen et al., 2010).

Our results suggest that when condition becomes more P limiting at higher rbCOD/P ratios, both energy and reducing power generation required for acetate uptake and PHB formation might shift from relying on both polyP hydrolysis and glycolysis pathway, to more enhancement and dependence on glycolysis in addition to partial/reverse TCA cycle. This is supported by The anaerobic glycogen utilization to polyP utilization ratio inside PAOs which increased nearly 20 times from 0.009 to 0.21 as influent rbCOD/P elevated from 20 to 50 (Table 2). This is also consistent with the increasing trend of glycogen_degradation_/HAc_uptake_ ratio with higher rbCOD/P ratio (Table 1). Furthermore, Figure 6 is a representation of the increase in the fraction of glycogen containing PAO cells (those containing both polyP and glycogen or polyP+glycogen+PHB) in the EBPR system under study and the corresponding increase in the PAO cells that contained PHV as the rbCOD/P ratio increases. Previously established EBPR models (SFigure 5) and current understanding (SFigure 6) assume that PHB can be formed via some form of TCA cycle or glycolysis or combination of both, however, PHV can mainly be formed through glycolysis pathway in combination with the succinate-propionate pathway (reductive branch) of TCA cycle or reverse TCA with glyoxylate shut combined with succinyl-CoA (daSilva et al., 2018) Thus, increase in the PHV containing PAO cells indicates the larger extent of employment of partial/split TCA pathways with increasing rbCOD/P ratio. Previous studies have often associated the increase in PHV content in the EBPR system with increase in the relative population abundance of GAOs since the latter mainly utilize glycolysis pathway and produce higher amount of PHV. However, our results revealed that the fraction of PAOs that contain PHV could vary as well with different operational loading conditions, irrelevant to the abundance of GAOs. Acevado et al (2012, 2017) showed that long-time P limited conditions enhance glycolytic pathway to supply energy deficit and shift towards more GAO-like metabolism. It was observed later by da Silva et al (2018) that when polyP is limited, reverse (split) TCA cycle is the most optimal pathway suggesting GAOs are operating reverse TCA cycle.

**Figure 6:**
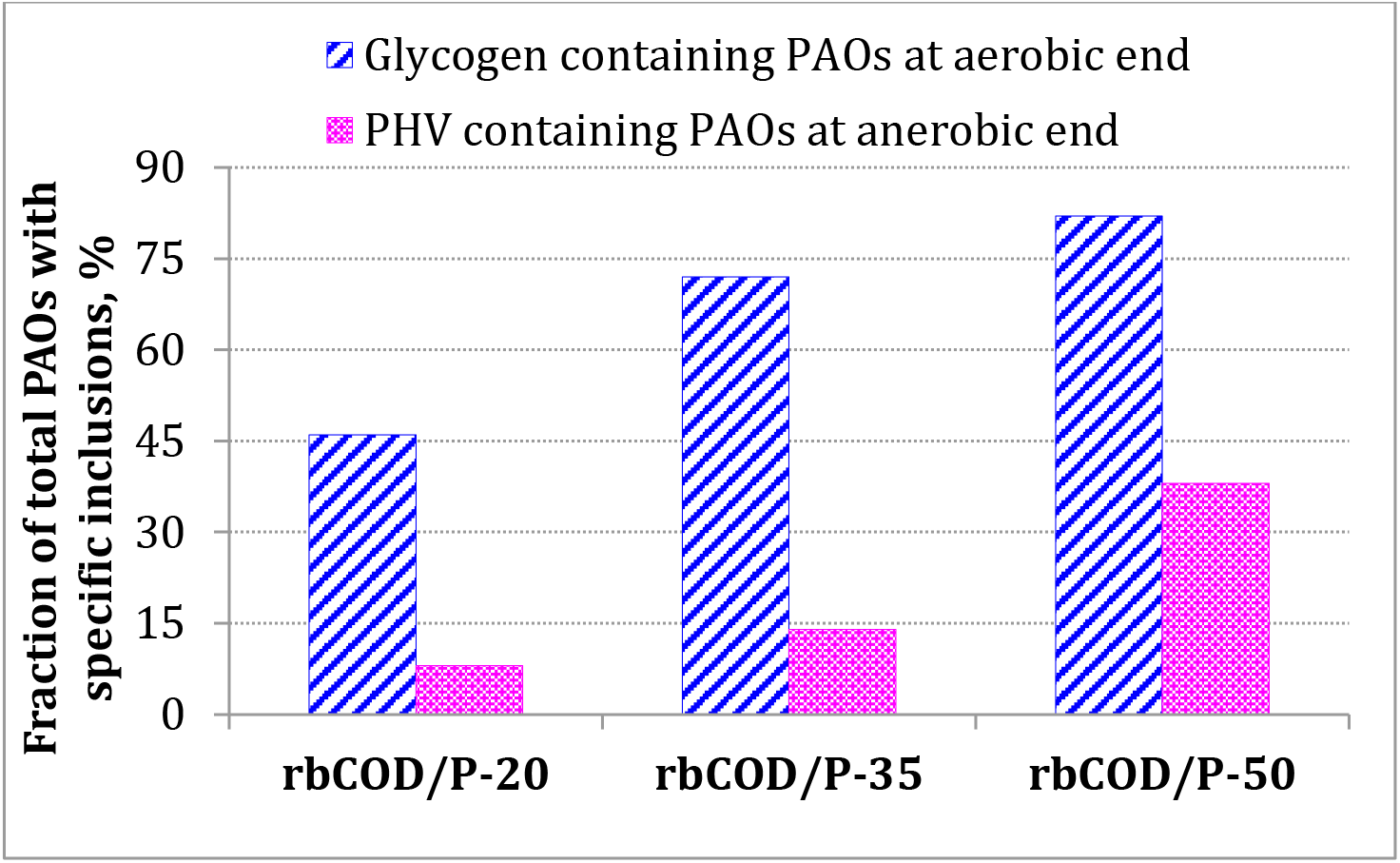
Distribution of fractions of PAOs (over total PAOs at respective phases) containing PHV at the end of anaerobic phase and glycogen at the end of aerobic phase in EBPR systems with different influent rbCOD/P ratios

It should be noted that for individual PAO cells in our study, the relative percentage of PHV content to total PHB+PHV never exceeded 8-12% (data not shown), which is consistent with the PHV content range reported previously with PAO-enriched (>70%) system (Oehmen et al., 2007). These results, for the first time, provided direct cellular and population level evidence for the possibility that at higher rbCOD/P loading condition when P becomes more limited, there were shifts in the involvement of different metabolic pathways in PAOs. Now, whether this was caused by phenotype changes or shifts in phylogenetic sub-populations that possess different pathways still requires confirmation via further investigation. Furthermore, it is also possible that some of these PAO cells might follow certain metabolism that does not involve the known and identifiable storage polymers.

## Conclusion

The lack of PAO isolates and the tools to monitor the quantities related to metabolic states of PAOs made it difficult to investigate the speculations that require observation from both phenotypic and phylogenetic aspects. The Raman microscopy method employed in this study helped us gaining further population and cellular level insights into the metabolic diversity among sub-populations within PAO group via tracking of intracellular polymeric inclusions in different metabolic stages of EBPR system, in contrast to the bulk measurements describing the EBPR process that are actually “apparent” sums of different PAOs groups with diverse metabolic pathways. To summarize, the following major conclusions could be drawn:

- Influent rbCOD/P ratio affected EBPR system stability and relative dominance/abundance of functionally relevant PAOs/GAOs
- Individual cellular level polymers level heterogeneity, distributions and stoichiometry ratios changed in both PAOs and GAOs populations in response to changes in influent carbon and P loading conditions.
- Intracellular polymeric stoichiometry evaluation within PAO populations enabled by Raman analysis elucidated the phenotypic elasticity in PAOs depending on influent loading conditions
- Quantification of differentiated PHB and glycogen inclusion levels in PAOs and GAOs showed that the influent rbCOD/P ratio increases, the excessive carbon beyond stoichiometric requirement for PAOs was diverted into GAOs.
- Population-specific evaluation of intracellular polymers evidenced that P limiting conditions at higher rbCOD/P ratios led to enhancement and increased reliance on glycolysis in addition to partial/reverse TCA cycle for anaerobic reducing power generation.

These findings provided new insights into the metabolic diversity of the functionally relevant populations in EBPR and population level formation for mechanistic EBPR model improvement and development. It also demonstrated the potential of application of Raman method as a powerful tool for the fundamental understanding of EBPR mechanism.

## Acknowledgement

This study was funded by scholarship to ITRI-NU (Industrial Translational Research Initiative at NU) from R&D of Veolia Water Solutions and Technologies, Sweden. Special thanks to Dr. Thomas Welander for his support. We greatly thank Professor Max Diem, Dr. Tatyana Chernenko and Ms. Evgenia Zuser in the department of Chemistry and Chemical Biology at Northeastern University for their advice and assistance in Raman microscopic analysis. We acknowledge Ms. Firouzeh Shiranian for her assistance with the monitoring of the lab-scale SBR-EBPR.

## Supporting Information

Additional supporting information has been included for clarification which are, STable 1: Details on oligonucleotide probes used for FISH in this study and their respective target groups; STable 2 : Performance of SBR-EBPR at different COD/P ratios studied; SFigure 1: Exemplary P release and uptake profiles of batch testing for sludge from 3 different EBPR SBRs systems fed with different influent COD/P ratios; SFigure 2: Intracellular polymer abundance distribution among individual GAO cells for PHB and glycogen polymers during batch testing for different COD/P ratios; SFigure 3: Effect of sample size on Single Cell Raman Spectra (SCRS); SFigure 4: Radial based function kernel and Eigen-decomposition used to investigate the sufficient sampling size; SFigure 5: Flow of 1 C mol Acetate and the resulting stoichiometry of intracellular polymers polyphosphate, PHB and glycogen in alternative anaerobic metabolic pathways in EBPR for PHB formation (earlier models); SFigure 6: Redox balance strategies for *Accumulibacter* under anaerobic conditions (recent understanding).

## References

Acevedo, B., Oehmen, A., Carvalho, G., Seco, A., Borrás, L., Barat, R., 2012. Metabolic shift of polyphosphate-accumulating organisms with different levels of polyphosphate storage. Water Research 46, 1889–1900.

Acevedo, B., Murgui, M., Borrás, L., Barat, R., 2017. New insights in the metabolic behavior of PAO under negligible poly-P reserves. Chemical Engineering Journal 311, 82–90.

APHA, 1998. Standard Methods for the Examination of water and wastewater. 20th ed.; American Public Health Association, Washington D.C.

Barnard, J., Shaw, A., Lindeke, D., 2005. In Using Alternative Parameters to Predict Success for Phosphorus Removal in WWTPs, 73rd Annual Water Environment Federation Technical Exposition and Conference, Washington DC, USA.

Broughton, A., Pratt, S., Shilton, A., 2008. Enhanced biological phosphorus removal for high strength wastewater with a low rbCOD:P ratio. Bioresource Technology 99, 1236–1241.

Cech, J. S., Hartman, P., 1993. Competition between Polyphosphate and Polysaccharide Accumulating Bacteria in Enhanced Biological Phosphate Removal Systems. Water Research 27 (7), 1219–1225.

Daims, H., Lucker, S., Wagner, M., 2006. daime, a novel image analysis program for microbial ecology and biofilm research. Environmental Microbiology 8(2), 200–213.

daSilva, L. G, Gamez, K. O., Gomes, J. C., Akkermans, K., Welles, L., Abbas, B., van Loosdrecht, M. C. M., Wahl, S. A., 2018. Revealing metablic flexibility of Candidus Accumulabacter phosphatis through rediz cofactor analysis and metabolic network modeling, bioRxiv Preprint.

Erdal, Z. K., 2002. The Biochemistry of EBPR: Role of Glycogen in Biological Phosphorus Removal and the Impact of the Operating Conditions on the Involvement of Glycogen. PhD Dissertation, VIrginia Tech, Blacksburg, VA, USA.

Filipe, C.D.M., Daigger, G.T., Grady, C.P.L., 2001. pH as a key factor in the competition between glycogen-accumulating organisms and phosphorus-accumulating organisms. Water Environ. Res. 73 (2), 223–232.

Gu, A. Z., Saunders, A.M., Neethling, J.B., Stensel, H.D., Blackall, L., 2005. Investigation of PAOs and GAOs and their effects on EBPR performance at full-scale wastewater treatment plants in US. Water Environment Research Foundation: Alexandria, Virginia, USA.

Gu, A. Z., Saunders, A., Neethling, J. B., Stensel, H. D., Blackall, L. L., 2008. Functionally Relevant Microorganisms to Enhanced Biological Phosphorus Removal Performance at Full-Scale Wastewater Treatment Plants in the United States. Water Environment Research 80 (8), 688–698.

Gu, A. Z., 2018. PAO metabolic Pathways, Water Environment Federation Report.

He, S., Gu, A. Z., McMahon, K. D., 2008. Progress toward understanding the distribution of Accumulibacter among full-scale enhanced biological phosphorus removal systems. Microbial Ecology 55, (2), 229–236.

He, S. M., Kunin, V., Haynes, M., Martin, H. G., Ivanova, N., Rohwer, F., Hugenholtz, P., McMahon, K. D., 2010. Metatranscriptomic array analysis of ‘Candidatus Accumulibacter phosphatis’-enriched enhanced biological phosphorus removal sludge. Environmental Microbiology 12 (5), 1205–1217.

Jenkins, D., Richards, M.G., Daigger, G.T., 2003. Manual on the Causes and Control of Activated Sludge Bulking and Foaming. 2nd edition, Lewis, London, UK.

Kong, Y.H., Beer, M., Rees, G.N., Seviour, R.J., 2002. Functional analysis of microbial communities in aerobic-anaerobic sequencing batch reactors fed with different phosphorus/carbon (P/C) ratios. Microbiology 148, 2299–2307.

Lanham, A. B., Oehmen, A., Saunders, A. M., Carvalho, G., Nielsen, P. H., Reis, M. A., 2013. Metabolic versatility in full-scale wastewater treatment plants performing enhanced biological phosphorus removal. Water research 47(19), 7032–7041.

Li, Y., Cope, H. A., Rahman, S. M., Li, G., Nielsen, P. H., Elfick A., Gu, A. Z., 2018. Toward Better Understanding of EBPR Systems via Linking Raman-Based Phenotypic Profiling with Phylogenetic Diversity Environmental Science & Technology 52 (15), 8596–8606.

Liu, W. T., Nakamura, K., Matsuo, T., Mino, T., 1997. Internal energy-based competition between polyphosphate- and glycogen-accumulating bacteria in biological phosphorus removal reactors - Effect of P/C feeding ratio. Water Research 31(6), 1430–1438.

Lopez-Vazquez, C.M., Oehmen, A., Hooijmans, C.M., Brdjanovic, D., Gijzen, H.J., Yuan, Z., van Loosdrecht, M.C.M., 2009. Modeling the PAO GAO competition: effects of carbon source, pH and temperature. Water Res. 43 (2), 450–462.

Majed, N., Matthaus, C., Diem, M. and Gu, A.Z., 2009. Evaluation of Intracellular Polyphosphate Dynamics in Enhanced Biological Phosphorus Removal Process Using Raman Microscopy. Environmental Science & Technology 43, (11), 5436–5442.

Majed, N., Gu, A. Z., 2010. Application of Raman Microscopy for Simultaneous and Quantitative Evaluation of Multiple Intracellular Polymers Dynamics Functionally Relevant to Enhanced Biological Phosphorus Removal Processes. Environmental Science & Technology 44 (22), 8601–8608.

Majed, N., Chernenko, T., Diem, M., Gu, A. Z., 2012. Identification of functionally relevant populations in enhanced biological phosphorus removal processes based on intracellular polymers profiles and insights into the metabolic diversity and heterogeneity. Environemntal Science & Technology 46, 5010–5017.

Martin, H. G., Ivanova, N., Kunin, V., Warnecke, F., Barry, K. W., McHardy, A. C., Yeates, C., He, S. M., Salamov, A. A., Szeto, E., Dalin, E., Putnam, N. H., Shapiro, H. J., Pangilinan, J. L., Rigoutsos, I., Kyrpides, N. C., Blackall, L. L., McMahon, K. D., Hugenholtz, P., 2006. Metagenomic analysis of two enhanced biological phosphorus removal (EBPR) sludge communities. Nature Biotechnology 24 (10), 1263–1269.

Muszyński, A., Miłobędzka, A., 2015. The effects of carbon/phosphorus ratio on polyphosphate- and glycogen-accumulating organisms in aerobic granular sludge. International Journal of Environmental Science and Technology 12(9), 3053–3060.

Neethling, J. B., Bakke, B., Benisch, M., Gu, A. Z., Stephens, S., Stensel, H. D., 2005. Factors Influencing the Reliability of Enhanced Biological Phosphorus Removal. report 01-CTS-3; Water Environment Research Foundation: Alexandria, Virginia, UK.

Oehmen, A., Lemos, P. C., Carvalho, G., Yuan, Z. G., Keller, J., Blackall, L. L., Reis, M. A. M., 2007. Advances in enhanced biological phosphorus removal: From micro to macro scale. Water Research 41, (11), 2271–2300.

Oehmen, A., Carvalho, G., Lopez-Vazquez, C. M., van Loosdrecht, M. C. M., Reis, M. A. M., 2010. Incorporating microbial ecology into the metabolic modelling of polyphosphate accumulating organisms and glycogen accumulating organisms. Water Research 44 (17), 4992–5004.

Randall, C.W., Barnard, J.L., Stensel, D.H., 1992. Design and Retrofit of Wastewater Treatment Plants for Biological Nutrient Removal. Technomic Publishing Co, Lancaster, PA.

Saunders, A. M., Oehmen, A., Blackall, L. L., Yuan, Z., Keller, J., 2003. The effect of GAOs (glycogen accumulating organisms) on anaerobic carbon requirements in full-scale Australian EBPR (enhanced biological phosphorus removal) plants. Water Science and Technology 47(11), 37–43.

Schuler, A. J., Jenkins, D., 2003. Enhanced biological phosphorus removal from wastewater by biomass with different phosphorus contents, part I: Experimental results and comparison with metabolic models. Water Environ Res 75(6), 485–498.

Schuler, A. J., 2005. Diversity matters: Dynamic simulation of distributed bacterial states in suspended growth biological waste-water treatment systems. Biotechnol. Bioeng. 91 (1), 62–74.

Smolders, G. J. F., Vandermeij, J., Vanloosdrecht, M. C. M., Heijnen, J. J., 1994. Model of the Anaerobic Metabolism of the Biological Phosphorus Removal Process - Stoichiometry and ph Influence. Biotechnology and Bioengineering 43(6), 461–470.

Stephens, H. M., Neethling, J. B., Benischer, M., Gu, A. Z., Stensel, H. D., 2004. Comprehensive analysis of full-scale enhanced biological phosphorus removal facilities., Water Environment Federation 77th Annual Conference and Exposition, New Orleans, LA, USA.

Streichan, M., Golecki, J.R., Schon, G., 1990. Polyphosphate-accumulating bacteria from sewage plant with different processes of biological phosphorus removal. FEMS Microbiol. Ecol. 73, 113–124.

Tchobanoglous, G., Burton, F.L., Stensel, H.D., 2003. In: Eddy, Ma (Ed.), Wastewater Engineering: Treatment and Reuse. McGraw-Hill, New York, NY.

Tetreault, M. J., Benedict A. H., Kaempfer, C., Borth, E. F., 1986. Biological Phosphorus Removal: A Technology Evaluation. J. Water Pollut. Control Fed., 58, 823–837.

Tu, Y., Schuler, A.J., 2013. Low acetate concentrations favor polyphosphate-accumulating organisms over glycogen accumulating organisms in enhanced biological phosphorus removal from wastewater. Environ. Sci. Technol. 47 (8), 3816–3824.

Whang, L. M., Park, J. K., 2006. Competition between polyphosphate- and glycogen-accumulating organisms in enhanced-biological-phosphorus-removal systems: Effect of temperature and sludge age. Water Environment Research 78, (1), 4–11.

Wilmes, P., Andersson, A. F., Lefsrud, M. G., Wexler, M., Shah, M., Zhang, B., Hettich, R. L., Bond, P. L., VerBerkmoes, N. C., Banfield, J. F., 2008. Community proteogenomics highlights microbial strain-variant protein expression within activated sludge performing enhanced biological phosphorus removal. Isme Journal 2(8), 853–864.

Zeng, R. J., Saunders, A. M., Yuan, Z. G., Blackall, L. L., Keller, J., 2003. Identification and comparison of aerobic and denitrifying polyphosphate-accumulating organisms.” Biotechnol Bioeng 83(2), 140–148.

Zhou, Y., Pijuan, M., Zeng, R. J., Yuan, Z. G., 2009. Involvement of the TCA cycle in the anaerobic metabolism of polyphosphate accumulating organisms (PAOs). Water Research 43(5), 1330–1340.

Zhou, Y., Pijuan, M., Oehmen, A., Yuan, Z. G., 2010. The source of reducing power in the anaerobic metabolism of polyphosphate accumulating organisms (PAOs) - a mini-review. Water Science and Technology 61(7), 1653–1662.

Zilles, J. L., Peccia, J., Kim, M. W., Hung, C. H., Noguera, D. R., 2002. Involvement of Rhodocyclus-related organisms in phosphorus removal in full-scale wastewater treatment plants. Applied and Environmental Microbiology 68 (6), 2763–2769.

