## Supplemental Data 1 for "Impact of Influent Carbon to Phosphorus Ratio on Performance and Phenotypic Dynamics in Enhanced Biological Phosphorus Removal (EBPR) System - Insights into Carbon Distribution, Intracellular Polymer Stoichiometry and Pathways Shifts"

**Journal**:  Water Research

**Title of Manuscript:**

Summary of SI:

| Figure no. | Figure Description | Page no. |
| --- | --- | --- |
| STable 1 | Details on oligonucleotide probes used for FISH (for Comparing PAOs/GAOs abundance with Raman measurements) and their respective target groups | S2 |
| STable 2 | Performance of SBR-EBPR for different COD/P ratios studied | S2 |
| SFigure1 | Exemplary P release and uptake profiles of batch testing for sludge from 3 different EBPR SBRs systems fed with different influent COD/P ratios. AN – Anaerobic phase; AE – Aerobic phase | S3 |
| SFigure 2 | Intracellular polymer abundance (intensity) distribution among GAOs cells for PHB, and glycogen quantities (as CCD counts) respectively, for overall duration of the phosphate release and uptake test for different COD/P ratios. *X* axis: fractions of cells that contained equal or higher amount of the specific level of polymeric inclusion at any given testing | S4 |
| SFigure 3 | Effect of sample size on Single Cell Raman Spectra (SCRS). Top: Quantitative relationship between sample size and diversity of SCRS as described in He et al. (2017); Bottom: Comparison of distribution of combinations of inclusions for different sample size (Different number of cells analyzed for same sample) | S5 |
| SFigure 4 | Radial based function kernel and Eigen-decomposition are used to investigate the sufficient sampling size to maintain the diversity of the respective microbial population in the lab-scale EBPR reactors. | S6 |
| SFigure 5 | Flow of 1 C mol Acetate and the resulting stoichiometry of intracellular polymers polyphosphate, PHB and glycogen in alternative anaerobic metabolic pathways in EBPR for PHB formation (earlier models) | S7 |
| SFigure 6 | Redox balance strategies for *Accumulibacter* under anaerobic conditions (recent understanding) | S8 |

**SUPPORTING INFORMATION**

**STable 1: Details on oligonucleotide probes available during study that were used for FISH (for Comparing PAOs/GAOs abundance with Raman measurements) and their respective target groups**

| **Probe** | **Sequence** | **Fluorescent labels** | **Specificity** | **% Formamide** | **Reference** |
| --- | --- | --- | --- | --- | --- |
| EUB338 | GCTGCCTCCCGTAGGAGT | FAM | Bacteria | 20 | [1] |
| PAO462 | CCGTCATCTACWCAGGGTATTAAC | CY3 | Most *Accumulibacter* | 35 | [2] |
| PAO651 | CCCTCTGCCAAACTCCAG | CY3 | Most  *Accumulibacter* | 35 | [2] |
| PAO846 | GTTAGCTACGGCACTAAAAGG | CY3 | Most *Accumulibacter* | 35 | [2] |
| Actino-221a | CGCAGGTCCATCCCAGAC | FAM | *Actinobacteria* | 35 | [3] |
| Actino-658a | TCCGGTCTCCCCTACCAT | FAM | *Actinobacteria* | 35 | [3] |
| GAOQ431 | TCCCCGCCTAAAGGGCTT | CY5 | Some *Competibacter* | 35 | [4] |
| GAOQ989 | TTCCCCGGATGTCAAGGC | CY5 | Some *Competibacter* | 35 | [4] |
| GB | CGATCCTCTAGCCCACT | FAM | Most *Competibacter* | 35 | [5] |
| DF988 | GATACGACGCCCATGTCAAGGG | CY5 | *Defluvicoccus* Cluster 2 | 35 | [6] |
| DF1020 | CCGGCCGAACCGACTCCC | CY5 | *Defluvicoccus* Cluster 2 | 35 | [6] |

**STable 2: Performance of SBR-EBPR at different influent COD/P ratios studied**

| **COD/P,**  **mg-COD/mg-P** | **Effluent P ±standard deviation, mg-P/L** | **P removal efficiency,**  **%** |
| --- | --- | --- |
| 20 | 0.43 ± 0.24 | 96.74 |
| 35 | 0.5 ± 0.25 | 94.99 |
| 50 | 1.82 ± 1.8 | 81.82 |

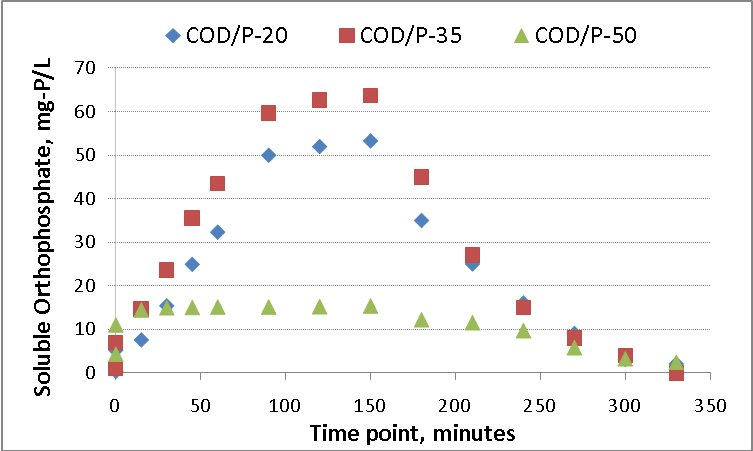

AN

AE

**SFigure1: Exemplary P release and uptake profiles of batch testing for sludge from 3 different EBPR SBRs systems fed with different influent COD/P ratios. AN – Anaerobic phase; AE – Aerobic phase**

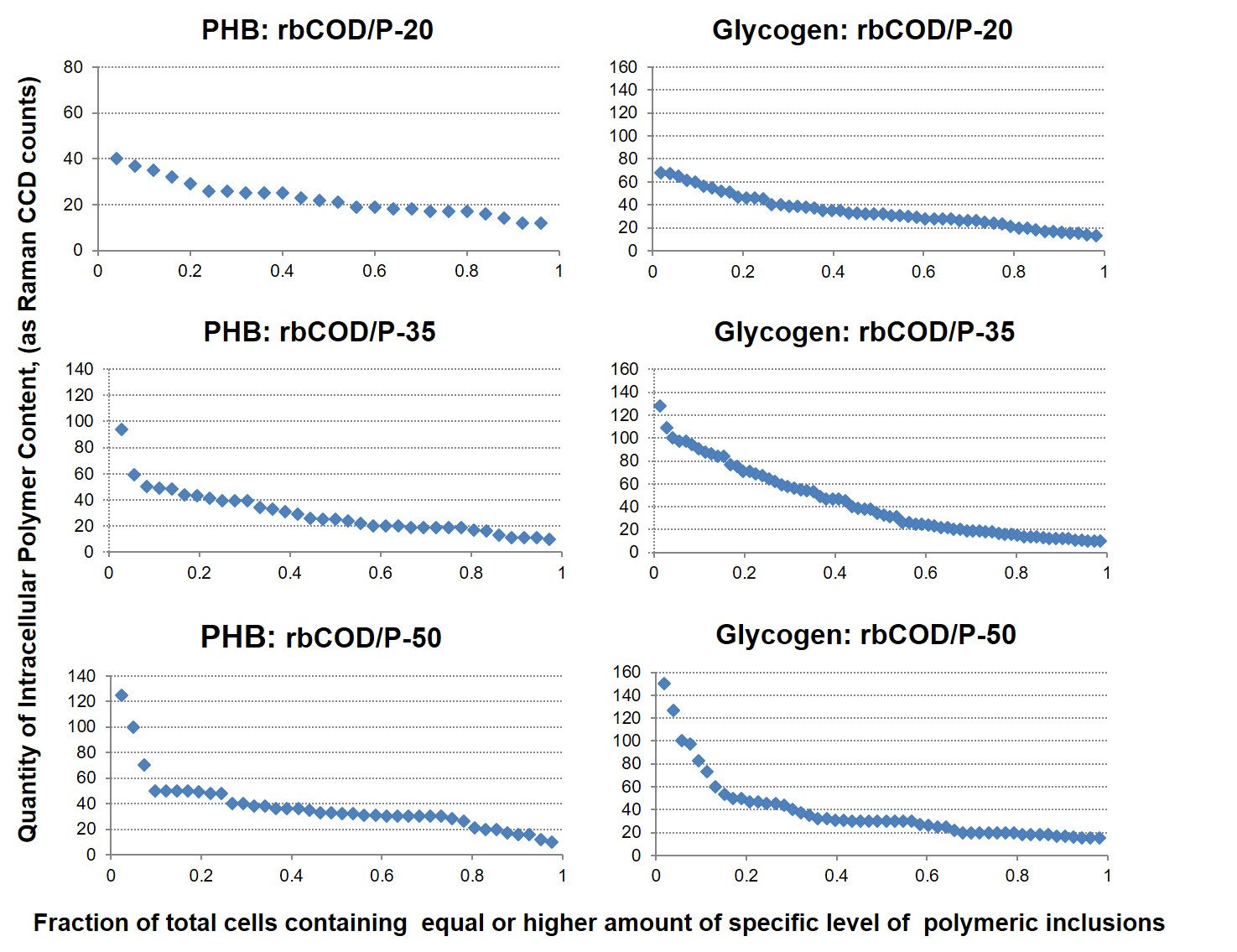

**SFigure 2: Intracellular polymer abundance (intensity) distribution among GAOs cells for PHB, and glycogen quantities (as CCD counts) respectively, for overall duration of the phosphate release and uptake test for different COD/P ratios. *X* axis: fractions of cells that contained equal or higher amount of the specific level of polymeric inclusion at any given testing.**

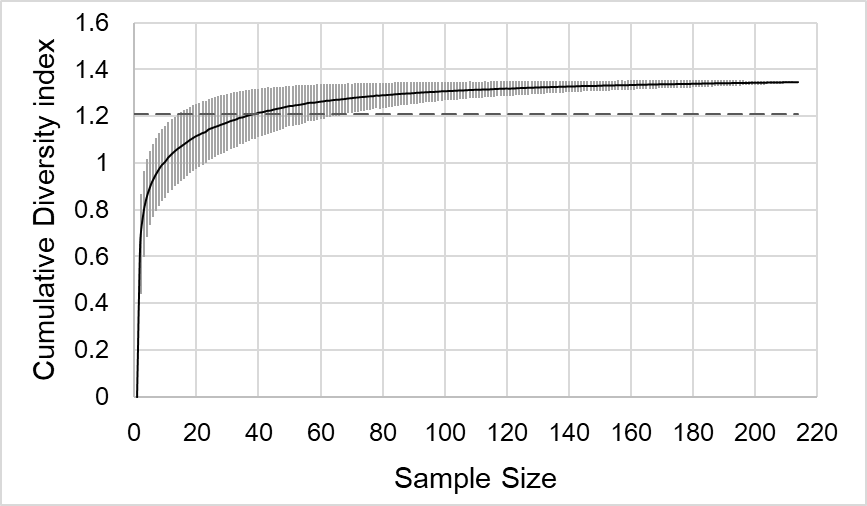

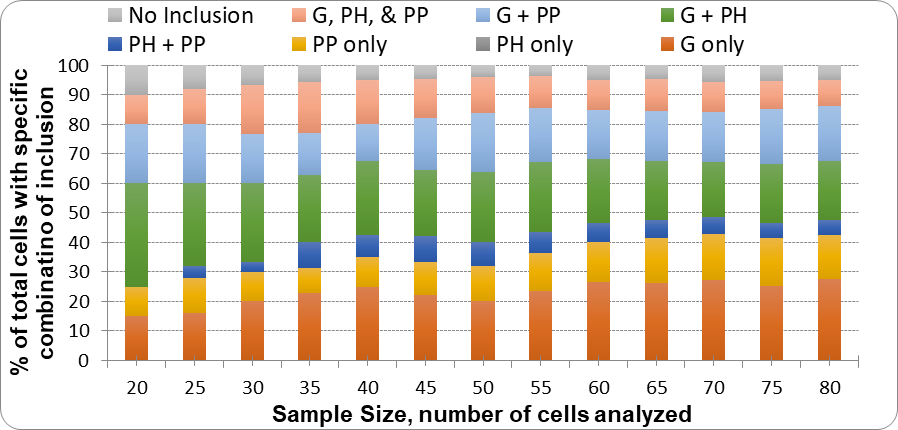

**SFigure 3: Effect of sample size on Single Cell Raman Spectra (SCRS). Top: Quantitative relationship between sample size and diversity of SCRS as described in He et al. (2017); Bottom: Comparison of distribution of combinations of inclusions for different sample size (Different number of cells analyzed for same sample).**

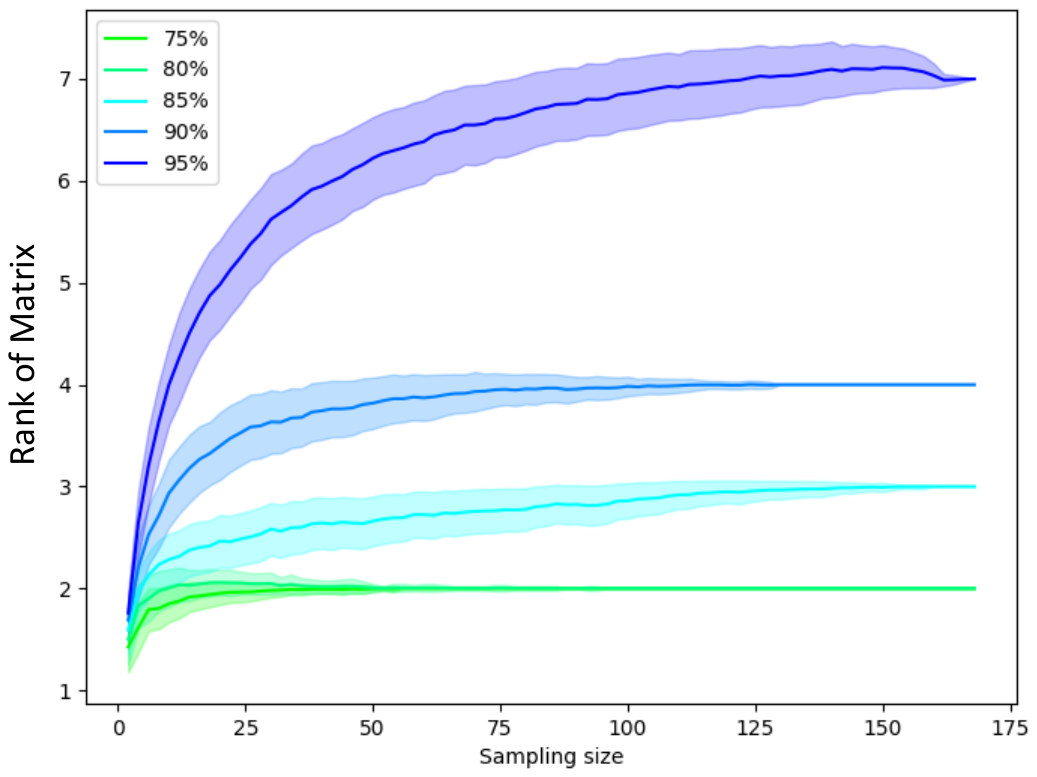

**SFigure 4: Radial based function kernel and Eigen-decomposition ^2^are used to investigate the sufficient sampling size to capture and maintain the diversity of the respective microbial population in the lab-scale EBPR reactors. This figure shows the mean and standard deviation (shaded area) of numbers of clusters found to preserve certain amount (%) of information form 500 random independent subsampling trials (Li et al 2018). 50 cells sampling size can capture approximately 85-90% of the “diversity” or information in the lab-scale EBPR reactor.**

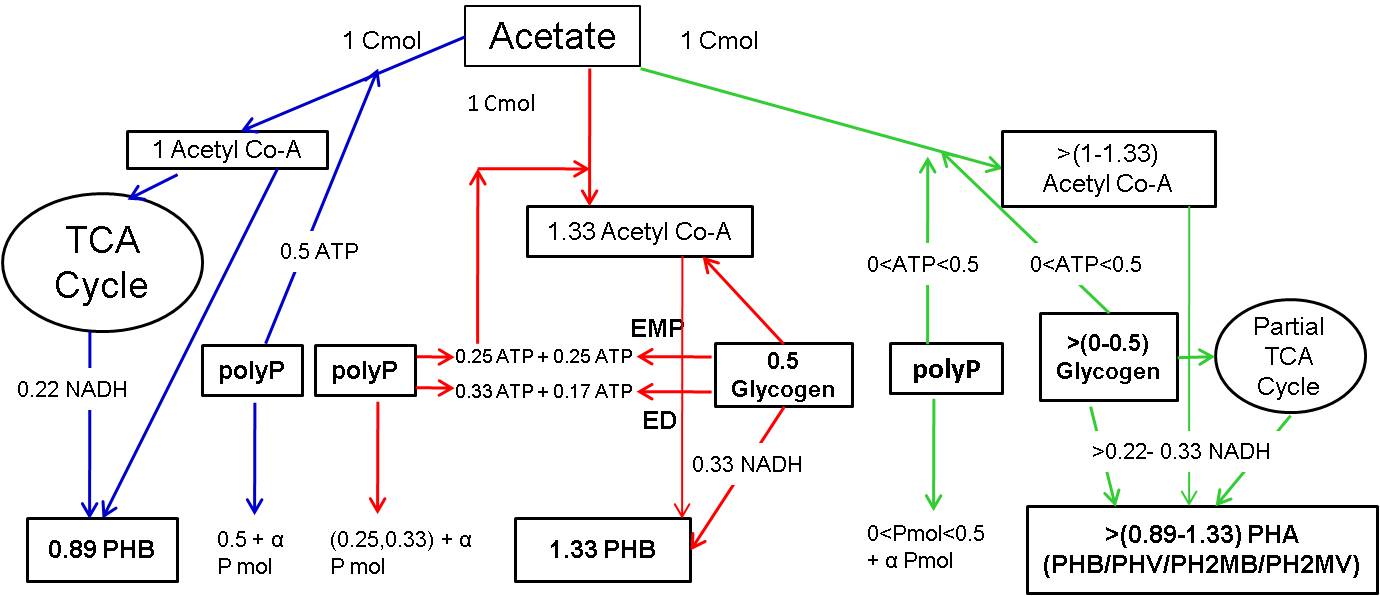

**SFigure 5: Flow of 1 C mol Acetate and the resulting stoichiometry of intracellular polymers polyphosphate, PHB and glycogen in alternative anaerobic metabolic pathways in EBPR for PHB formation. Blue arrow represents TCA cycle only, Red arrows showing glycolysis only and green arrows represent combined glycolysis + partial TCA cycle pathways. α represents the amount of energy required to transport 1 C-mol of acetate (Combination of Comeau model (TCA only), Mino/adapted Mino model (glycolysis only) and Perreira, Yagci, Hesselman model (glycolysis + partial TCA))**

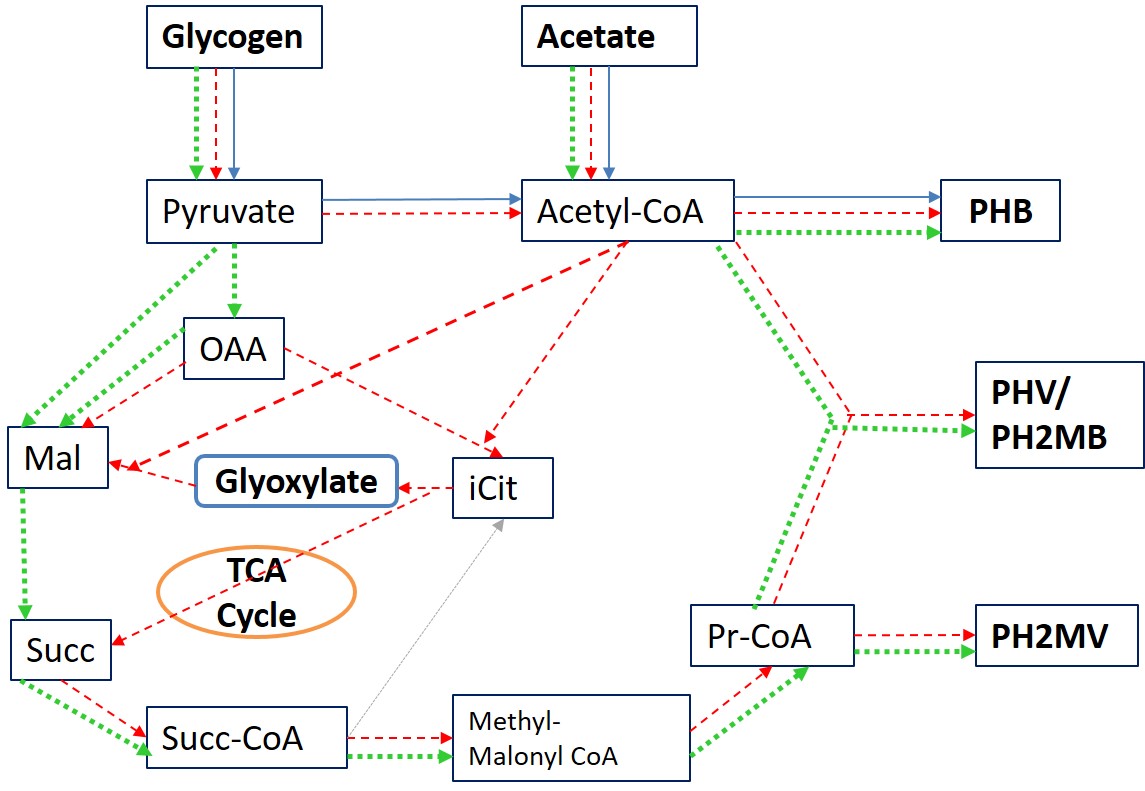

**SFigure 6: Redox balance strategies for *Accumulibacter* under anaerobic conditions according to daSilva et al (2018). Blue arrows represent glycogen degradation for acetate reduction to PHB when stoichiometric amount is degraded; Red dotted arrows represent glycogen degradation in addition to glyoxylate shunt for PHB and PHA generation when glycogen is limited; green dotted arrows represent glycogen degradation in addition to reductive TCA branch when glycogen is abundant.**

References:

Amann, R. I., B. J. Binder, R. J. Olson, S. W. Chisholm, R. Devereux, and D. A. Stahl, 1990. Combination of 16S rRNA-targeted oligonucleotide probes with flow cytometry for analyzing mixed microbial populations. Appl. Environ. Microbiol. 56, 1919-1925.

Crocetti, G. R.; Hugenholtz, P.; Bond, P. L.; Schuler, A.; Keller, J.; Jenkins, D.; Blackall, L. L., 2000. Identification of polyphosphate-accumulating organisms and design of 16S rRNA-directed probes for their detection and quantitation. Applied and Environmental Microbiology. 66, (3), 1175-1182.

Crocetti, G. R.; Banfield, J. F.; Keller, J.; Bond, P. L.; Blackall, L. L., 2002. Glycogen-accumulating organisms in laboratory-scale and full-scale wastewater treatment processes. Microbiology-Sgm. 148, 3353-3364.

Kong, Y. H.; Nielsen, J. L.; Nielsen, P. H., 2005. Identity and ecophysiology of uncultured actinobacterial polyphosphate-accumulating organisms in full-scale enhanced biological phosphorus removal plants. Applied and Environmental Microbiology. 71, (7), 4076-4085.

Kong, Y. H.; Ong, S. L.; Ng, W. J.; Liu, W. T., 2002. Diversity and distribution of a deeply branched novel proteobacterial group found in anaerobic-aerobic activated sludge processes. Environmental Microbiology. 4 (11), 753-757.

Meyer, R. L.; Saunders, A. M.; Blackall, L. L., 2006. Putative glycogen-accumulating organisms belonging to the Alphaproteobacteria identified through rRNA-based stable isotope probing. Microbiology-Sgm., 152, 419-429.
